# Spp1 secreted by macrophages impairs osteogenic ability of Ctsk^+^Fmod^+^ periosteal cells in jaw bone

**DOI:** 10.1101/2023.05.23.541910

**Authors:** Zumu Yi, Yeyu Liu, Jing Wang, Chen Hu, Yi Man

**Author notes:** Corresponding author. Yi Man, DDS, Ph.D, Professor and Chair, Department of Oral Implantology, West China Hospital of Stomatology, Sichuan University, 14#, 3rd section, Renmin South Road, Chengdu 610041, China. Chen Hu, Ph.D, Department of Oral Implantology, West China Hospital of Stomatology, Sichuan University, 14#, 3rd section, Renmin South Road, Chengdu 610041, China. These authors contributed equally to this work (Zumu Yi, Yeyu Liu, Jing Wang). **Disclosure of Interest** The authors state no conflict of interest.

## Abstract

Periosteum, which covers the surface of most bones, mediates bone regeneration through endochondral ossification during fracture repair and intramembranous ossification under steady state. Periosteal cells (PCs) of jaw bones are different from those of long bones in phenotypic characteristics and functions. So far, the role of periosteum in jaw bones during bone grafting remain unclarified. Here we propose a subperiosteal bone grafting model based on the clinical procedures. By integrating single-cell RNA sequencing (scRNA-seq) and spatial transcriptomic (ST), we found a functional Ctsk^+^Fmod^+^ subset of PCs in jaw bones. The Ctsk^+^Fmod^+^ PCs had the potential of multi-directional differentiation. Furthermore, Spp1 secreted by macrophages could impair the osteogenic capacity of Ctsk^+^Fmod^+^ PCs, which could be partly rescued by blocking Spp1. The identification of this Ctsk^+^Fmod^+^ subclusters, which shows osteoprogenitor characteristics and close interaction with macrophages, reveals the heterogeneity of periosteal cells in jaw bones, and may provide target of intervention to improve osteogenesis during bone augmentation surgery.

## Introduction

Periosteum, a complex and orderly connective tissue envelope with abundant blood supply, covers the surface of most bones(*1*). Derived from perichondrium, the final periosteum can be generally divided into inner and outer layers(*2*). It contains different types of cells, ranging from mesenchymal stromal cells (MSCs) to myeloid lineage cells (MCs), osteogenic cells and fibroblasts(*3, 4*). Periosteal cells (PCs) are highly heterogeneous. Cathepsin K^+^(Ctsk^+^) periosteal stem cells(PSCs) mediate bone regeneration through endochondral ossification during fracture repair and intramembranous ossification under steady state(*5*). Mx1 and αSMA can label a subpopulation of PSCs, which are required for injury repair(*6*). Sox9-expressing PSCs initiate cartilage callus formation via giving rise to skeletal cells(*7*). Markers like Platelet-derived growth factor receptor α(PDGFRα), Grem1, Gli1, Nestin (Nes) have also been used to identify PSCs(*8–10*). However, due to the cellular heterogeneity and functional divergence within PSCs, the determination of these progenitor cells remains a challenge.

Accumulating evidence demonstrates that PSCs of jaw bones display unique biological characteristics due to the distinct differences in developmental, mechanical, or homeostatic properties of long bones and jaw bones(*11–13*). The local ablation of Ctsk^+^Ly6a^+^ jawbone PSCs delayed fracture repair(*11*). However, so far studies on PSCs of jaw bones are still scarce. The functional and phenotypic characteristics of PSCs in jaw bones, and how PSCs react during alveolar bone grafting remain unclarified.

In addition, the immune microenvironment is more active in alveolar bone and the macrophages most actively interactwith MSCs(*14*). Macrophages can regulate tissue regeneration in specific microenvironment(*15*). Tartrate-resistant acid phosphatase positive (TRAP^+^) macrophages can recruit Nes^+^ and Leptin receptor^+^ PSCs for periosteal bone formation by secreting platelet-derived growth factor-BB (PDGF-BB) (*16*). The CD68^+^F4/80^+^ macrophages of periosteum express and activate transforming growth factor β(TGF-β1) to recruit PSCs(*3*). Recently, the SPP1^hi^ macrophages has gradually attracted the interest of researchers. Studies demonstrated SPP1^hi^ macrophages contribute significantly to fibrosis in different tissues and organs(*17*). Nevertheless, whether these cells act on jaw bone PCs remains to be explored. It’s important to understand how macrophages are involved in the regulation of PCs in jaw bone periosteal microenvironment.

To identify the PCs subpopulations in jaw bones and the effect of macrophage on the cell function of PCs, we integrated high-throughput single-cell RNA sequencing (scRNA-seq) and spatial transcriptomic (ST) to delineate the characteristics of jaw bone’s periosteal microenvironment. Clinicians have modified the guided bone regeneration (GBR) procedures by using intact periosteum to cover implanted bone grafts, on account of the regenerative potentials of periosteum (*18, 19*). Based on that, we propose a subperiosteal bone grafting model. In this study, we report a functional Ctsk^+^Fmod^+^ subset of PCs in the jaw bones. This subpopulation has the ability to differentiate into a variety of cells, such as osteoblasts, chondroblasts and adipocytes. When bone graft substitutes are implanted under the periosteum of the jaw bones, the anti-inflammatory macrophage (AIM), pro-inflammatory macrophages (PIM) and Ctsk^+^Fmod^+^ PCs would migrate to the vicinity of the materials. The AIM and PIM secrete Spp1, which impairs the osteogenic ability of Ctsk^+^Fmod^+^ PCs. Blocking Spp1 could increase the osteogenic potential of Ctsk^+^Fmod^+^ PCs and promoted osteogenesis in the jaw bone periosteal microenvironment.

## Results

### Establishment of subperiosteal bone grafting model

To evaluate hard tissues changes in GBR with intact periosteum, 40 patients from 2020 to 2021 were analyzed(*20*). Histological analysis of samples taken 6 months after surgery suggested that there was new bone formation underlying the periosteum. Cone-beam computed tomography (CBCT) images demonstrated that comparable and acceptable results of horizontal bone augmentation can be achieved with the technique of GBR with intact periosteum.

To further understand the role and characteristics of jaw bone periosteal cells, the subperiosteal bone grafting model was established inspired by the clinical operation and literature report(*21*)(fig. 1A). To ensure the integrity of periosteum, we evaluated evaluated the attachment of the periosteum after being elevated. It was noted that the periosteum in the untreated group was attached to the hard palate, while the periosteum in the operated group was completely turned up from the bony surface(fig. S1A). To verify whether the subperiosteal bone grafting model could induce the de novo bone formation as human, we harvested samples at different time points for analysis. More blood cells and fibrin networks could be found in Bio group on day 3(fig. S1B). At post-surgical day 7, osteoblasts and osteoid deposition were apparent near the bone graft(fig. 1B). Notably, the regular periosteal structure was lost in the Bio group, suggesting that the periosteum was undergoing remodeling. New bone formation occurred at day 14 and there was obvious new bone formation in intimate contact with bone grafts in Bio group at day 28(fig. S1C, fig. 1C). In control group, periosteal thickening occurred after surgery and there was no significant new bone formation. Immunofluorescent staining showed that osteogenesis-related markers (ALP, OCN and OPN) were expressed around the bone grafts, while these markers were mainly expressed in the alveolar bone in Ctrl group(fig. S1D). Taken together, these data showed that implantation of bone grafts under the periosteum could induce vertical bone regeneration in the jaw bone and the subperiosteal bone grafting model was successfully established.

**Fig. 1.**
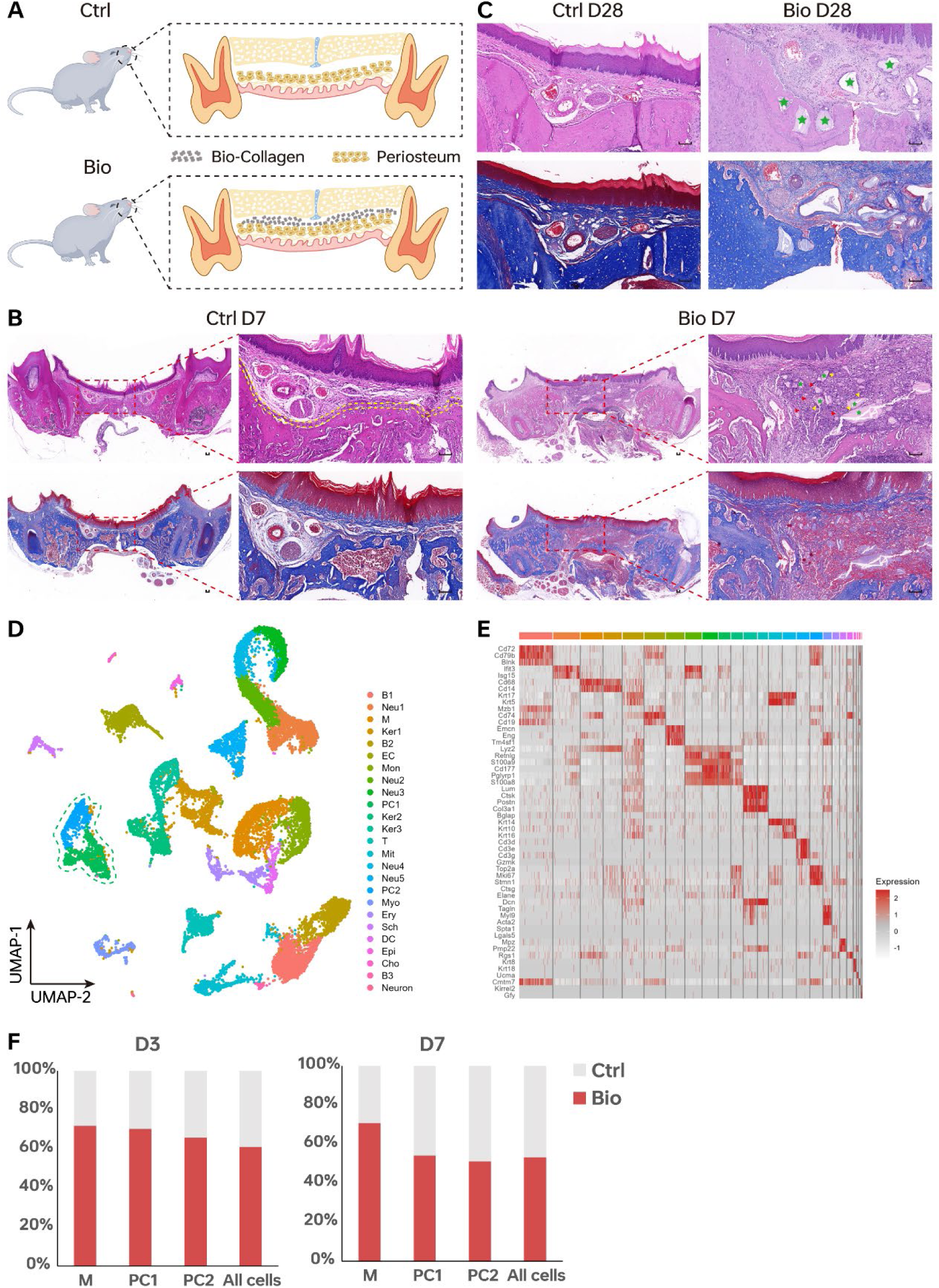
The establishment of a rat subperiosteal bone grafting model and Single-cell landscape of Bio and Ctrl group at different timepoints. (A) Schematic diagram of rat subperiosteal bone grafting model. Representative H&E and Masson staining images of Ctrl and Bio group on D7(B) and D28(C). Yellow dotted line, periosteum; Yellow arrows, osteoblasts; Red arrows, osteoid deposition; Green stars, bone grafts. Scale bar, 100μm. (D) Uniform Manifold Approximation and Projection (UMAP) plots of 19377 cells from different groups, showing 25 clusters. Green dotted line, PC-related clusters. (E) Heatmap showed top marker genes for each cluster. (F) Bar plot of proportion of 3 selected clusters in different group at different timepoints.

### Single-cell sequencing identifies major cell composition of overlying soft tissue on rat jaw bone

In order to identify the PCs subpopulations in jaw bones, three fresh samples from the surgical area were harvested per group (Bio and Ctrl) at different timepoints (Day3 and Day7) (fig. S2A). We processed these samples into single-cell suspensions and profiled them with scRNA-seq. Our dataset get 19377 cells (Day3 Bio[n=6254 cells], Day3 Ctrl[n=3999 cells], Day7 Bio[n=4849 cells] and Day7 Ctrl[n=4275 cells]) after quality control(fig. S2B-C). we used CCA to correct the batch effect (fig. S2C). We got 25 clusters and annotated each cluster with their respective markers (fig. 1, D-E;Table S1). Identified periosteal cell (PC) (Ctsk, Postn and Col3a1), macrophage (Cd68, Cd14) and monocyte-macrophage (Fcnb, Cd14) marker genes were cell type specific(fig. S2D). We compared the cells proportion of each group at different times and found that the periosteum cell population (PC1 and PC2) increased on day 3 due to bone graft substitute implantation, and there was no significant difference between the two groups on day 7. At the meantime, the macrophages had a higher cell proportion at all time points in Bio group (fig. 1F, fig. S2E).

### Heterogeneity analysis within the PCs and identification of Ctsk^+^Fmod^+^ PCs in rat jaw bone

We extracted cells defined as PCs and sub-divided them into 12 clusters. Among them, we defined PCs-related cells, including periosteal cell 1 (Ctsk, Fmod, Tnmd), periosteal cell 2 (Ctsk, Prrx1, Pdgf α), chondrocyte (Sox6, Ucma), pre-osteoblast (Alpl, Ostn), fibroblast (Acta2, Tagln), osteoblast (Alpl, Bglap) and Nes^+^ periosteal cell (Nes, Dlx5) based on marker gene profiles published before(fig. 2, A-B)(*5, 8, 10, 22*). We also found a group of periosteal stromal cells containing fibroblast/stromal cells and Ecm1^+^ fibroblasts(fig. S3A). To analyze the function of each subpopulation, we obtained top marker genes for each cluster and conducted GO analysis. GO analysis showed that parts of subclusters of PCs were related to ossification, connective tissue development and mesenchymal cell differentiation, especially periosteal cell 1(fig. 2C). Periosteal stromal cells were related to extracellular matrix and structure organization(fig. S3B). Moreover, periosteal cell 1 highly expressed stemness-related markers (Ctsk, Pdgfα and Prrx1). Therefore, we speculated that periosteal cell 1 was a population of cells having differentiation potential (Ctsk^+^Fmod^+^ PCs)(fig. 2C; fig. S3C).

**Fig. 2.**
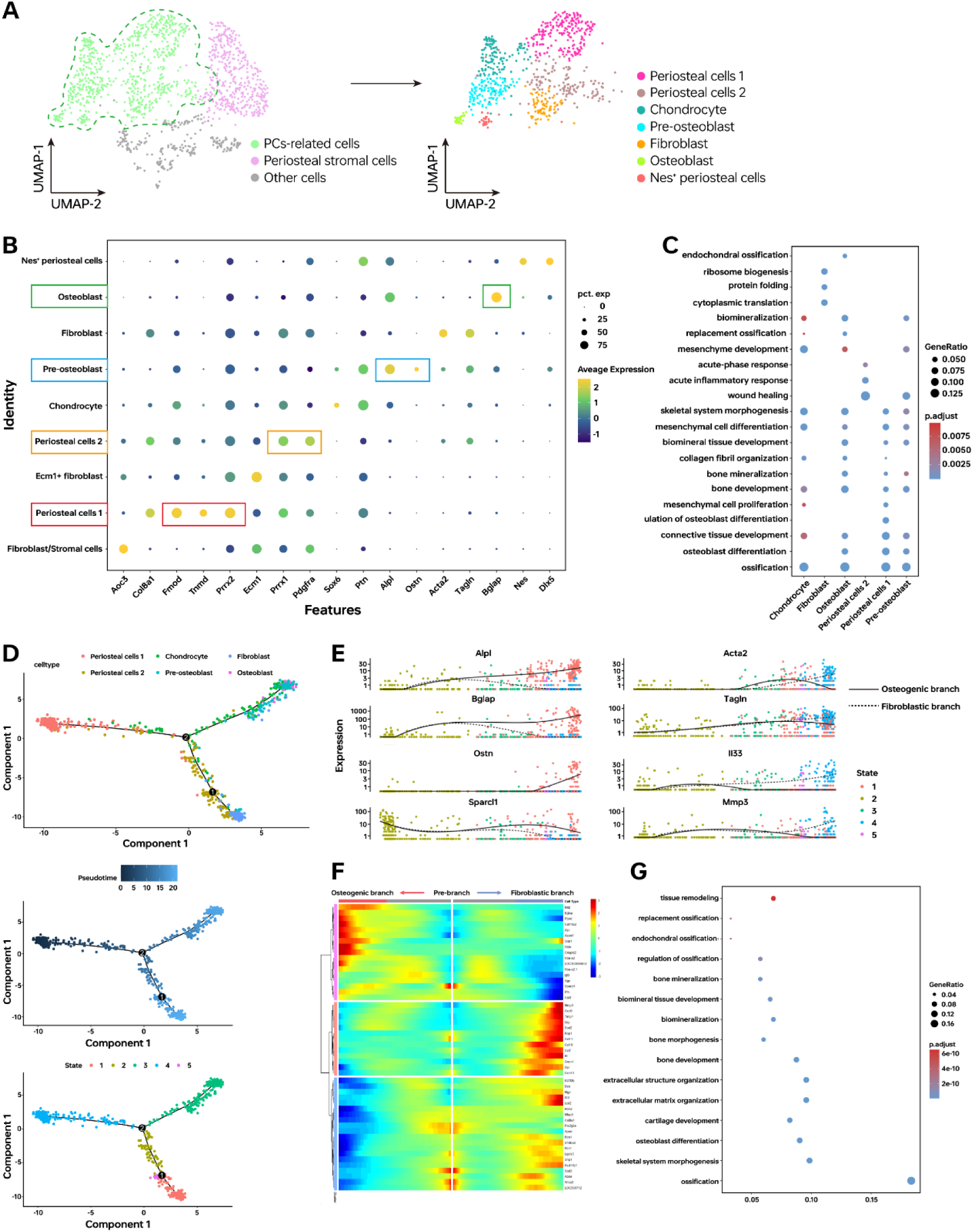
Periosteal cell identification and characterization in Jaw bone. (A) UMAP plots showing subclustering results of PCs. (B) Dot plots showing expression percent and average expression of markers in subtypes of PCs. (C) GO analysis of subclusters of PSCs-related cells. (D)Trajectory reconstruction of PCs based on a pseudo-temporal order from left to right; color-coding by celltype. (E) Relative gene expression dynamics of genes in cells on osteogenic and fibroblastic trajectory. (F) Gene expression heatmap of branch 2 in a pseudo-temporal order. Osteogenic and Fibroblastic trajectories are shown on the left and right, respectively. (G) GO analysis of differentially expressed genes (DEGs) in cells comparing the osteogenic branch with fibroblastic branch.

Similar to the previous analysis, the number of all periosteal subsets (except Nes^+^ periosteal cell) in the Bio group was higher than that in the Ctrl group on day 3(fig. S3D). This trend disappeared on day7, but a higher proportion of fibroblasts and pre-osteoblasts remained(fig. S3D). Then we compared the differentially expressed genes (DEGs) between different groups. On day3, osteogenic related activities, such as ossification and cartilage development, were stronger in the Bio group(fig. S3E). However, when it turned to day7, immune-related functions were enhanced and leukocytes, especially granulocytes, were recruited(fig. S3F). Compared with day3, osteogenic related activities were decreased in day7 Bio group.

Overall, our data reveal a complex landscape of PCs in Rat Jaw bone. PCs contained multiple different cell subtypes with different functions, among which Ctsk^+^Fmod^+^ subset might be a population of cells having differentiation potential. When the periosteum of jaw bone was stimulated by external stimuli (like bone graft substitute), it would activate the periosteum. The intervention of immune cells might have an important influence on this process.

### Ctsk^+^Fmod^+^ PCs underwent osteogenic differentiation

To generate a pseudo-temporal map of differentiation trajectories of PCs subtypes, we used Monocle2, an algorithm to reconstruct biological processes according to transcriptional similarity. Trajectory analyses demonstrated periosteal cell 1(Ctsk^+^Fmod^+^ PCs) located at upstream and the Ctsk^+^Fmod^+^ PCs bifurcated into two diverse branches including fibroblast and osteoblast, respectively (fig. 2D). This trend was consistent with changes in cell density(fig. S3G). The cells in one terminal expressed genes associated with bone formation (like *Alp*, *Bglap*, *Ostn*) while another maintained high expression levels of fibroblastic genes(fig. 2E).

Differential expression per branch was analyzed to get insights into the gene expression changes along the trajectory. In comparison with the fibroblastic branch, cells in osteogenic branch expressed higher levels of genes related to osteogenesis, which were enriched for the GO terms such as “Ossification”, “skeletal system morphogenesis”, “osteoblast differentiation” and “bone development” (fig2.F-G). Taken together, these gene expression patterns characterized the potential role of Ctsk^+^Fmod^+^ PCs. When Ctsk^+^Fmod^+^ PCs were stimulated, it might contribute to the generation of osteogenic cells, with the activation of genes related to osteogenesis.

### Characteristics of macrophages in different groups

Macrophages have been reported to regulate tissue regeneration and there are close interactions between macrophages and PCs(*3, 23*). To explore the heterogeneity of macrophages, the macrophages, monocyte-macrophages and dendritic cells were extracted and then reclustered into 9 subpopulations(figS4A). According to known markers(*24, 25*), we defined pro-inflammatory macrophages(PIM)(Arg1,Mmp12,Tgm2), monocytes 1&2(Mono 1&2)(Vcan,Cd14,Camp), anti-inflammatory macrophages(AIM)(Mrc1,Folr1,C1qc), cycling cell 1&2(Top2a, Stmn1, Tpx2), dendritic cells 1&2(DC 1&2)(Cd74,Cytip) and osteoclast(Ctsk, Slc9b2)(figS4A-B). Similarly, we compared the cells proportion of macrophage subtypes at different times. In the Bio group, the number of PIM and AIM was more than Ctrl group, suggesting that the implantation of bone graft substitute recruited more macrophages (fig. S4C). To sum up, bone graft substitute implantation recruited more AIM and PIM to the periosteum, indicating differences in immune micro-environment of Bio and Ctrl groups.

### Co-localiztion of PC and macrophage subclusters revealed by spatial transcriptomics

To further assess the spatial characteristic of PCs and macrophages, we performed spatial transcriptomics (ST) with tissue sections from different groups at Day3 and Day7(fig. S5A). Transcriptomics from 1152 and 1033 spots were obtained at a median depth of 1694 and 2417 genes/spot(fig. S5B). The morphological structure of the tissues was easily discerned histologically(fig. 3A). Then ST and scRNA-seq gene expression profiles were integrated using the Addmodulescore function in Seurat(*26*). We used top50 differentially expressed marker genes of subtypes of PCs and macrophages to score spots gene expression profiles. Scoring spots with genes of periosteal cell 1 from scRNA-seq showed a cluster highly associated with periosteal cell 1 in ST (Cluster 6) (fig. S5C-D). According to the integration of scRNA-seq and ST at different time points, subclusters of periosteal cells (especially Ctsk^+^ Fmod^+^ PCs) in the Bio group were mainly located above the bone graft substitute at Day3(fig. 3B). When it turned to Day7, Ctsk^+^ Fmod^+^ PCs and periosteal cell 2 migrated around the bone graft substitute, while the two subclusters of PCs remained in situ in the periosteum in the Ctrl group(fig. 3C). Notably, both osteoblast-related cells (pre-osteoblast and osteoblast) and fibroblast-related cells(fibroblast) were present around the bone graft substitute on day 7, suggesting that fibroblastic and osteogenic activity co-existed(fig. 3C).

**Fig. 3.**
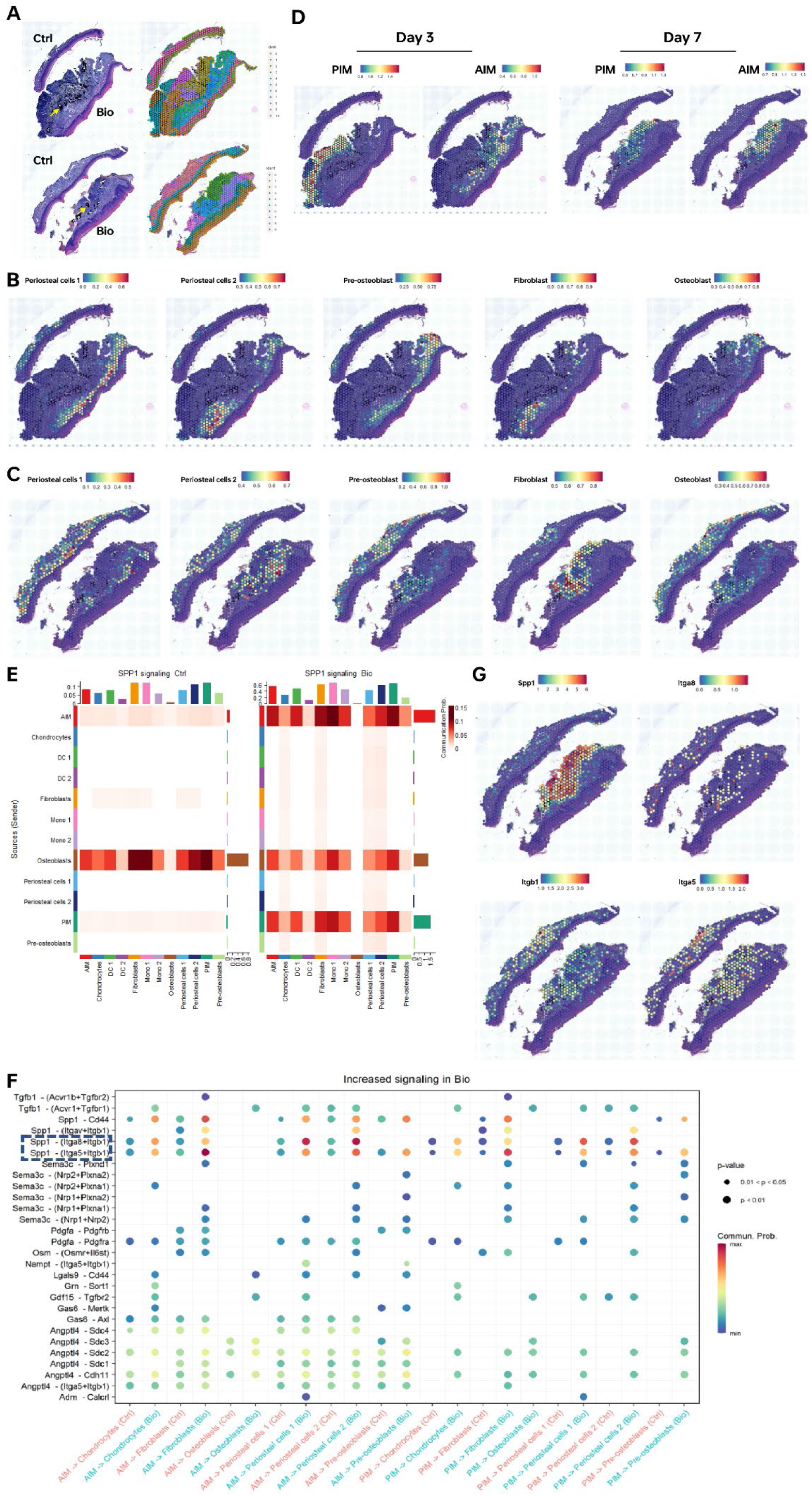
Spatial localization of PCs and macrophages revealed by spatial transcriptomics and mapping cellular crosstalk between PCs and macrophages. (A) HE staining of tissue sections and unbiased clustering of ST spots (Yellow arrow: bone graft substitute) (left). Spatial feature plots showing defined subtypes of PCs distribution in tissue sections at Day3(B) and Day7(C), as well as the distribution of macrophage subclusters at different timepoints(D). (E) Heatmap of SPP1 signaling pathways in Bio and Ctrl group. (F) Network plot showing the ligand-receptors analysis of Bio and Ctrl group. (G) ST showed select ligands and receptors expression.

Next, we analyzed the spatial localization of PIM and AIM by integrating scRNA-seq and ST. The spatial localization of PIM and AIM was consistent with PCs subtypes. PIM and AIM gradually migrated around the bone graft substitute over time (fig. 3D; fig. S5E). Score spots in each cluster with PCs subtype (Ctsk^+^ Fmod^+^ PCs and periosteal cell 2) or macrophages subtype (PIM and AIM) signatures co-localized in the same spot at Day7(fig3. C-D). As the cells were more likely to interact with nearby cells(*27*), the macrophage subtypes(PIM and AIM) may communicate with Ctsk^+^ Fmod^+^ PCs and affect the function of them.

Together, when bone graft substitute was implanted, Ctsk^+^ Fmod^+^ PCs were activated and gradually migrated towards the material to perform their applied functions. At the same time, PIM and AIM also migrated to the material and co-localized with PCs subtypes (Ctsk^+^ Fmod^+^ PCs and periosteal cell 2).

### PIM and AIM interacted with Ctsk^+^Fmod^+^ PCs via SPP1 signalling

Due to the close spatial location of Ctsk^+^Fmod^+^ PCs and macrophage subsets (PIM and AIM) at the Day7, it was suggested that they might have potential cellular communication. To further identify the key mediators of PCs and macrophages, we applied Cellchat(*28*) to investigate the cell–cell interaction among them. We compared the interaction strength (information flow) between Bio and Ctrl group(fig. S6A). The top increased signaling pathways colored blue were enriched in Bio group, including SPP1 and TGF-β signaling. The net plot showed that AIM radiated the most interaction strength among all cell types(fig. S6B). Next, we compared significant pathways between different clusters in Bio and Ctrl group, among which the SPP1 signaling pathway was one of the most prominent (fig. S6C-D). Further exploration of the SPP1 signaling pathway indicated that PIM and AIM sent strong signals, which were received by subsets of periosteal cells (Ctsk^+^Fmod^+^ PCs, periosteal cell 2 and fibroblast) in Bio group (fig. 3E). Detailed interaction network between cells showed the ligands and receptors, among which Spp1-(Itga8 + Itgb1) and Spp1-(Itga5 + Itgb1) signaling pathways were most prominent in Bio group(fig. 3F). The analysis of Spp1 expression in macrophages of different groups showed that *Spp1* was mainly expressed in Bio group(fig. S6E). PIM and AIM were the macrophage subsets that predominantly express *Spp1*(fig. S6E). We further examined the expression positions of these ligands in space with ST. Spatial plots showed ligand (Spp1) and receptors (Itga8+Itgb1;Itga5+Itbg1) preferentially aggregated around bone graft substitute(fig. 3G). The results of Cellchat and ST indicated that PIM and AIM could communicate with PCs including Ctsk^+^Fmod^+^ PCs, thus triggering downstream biological effects.

In summary, the macrophages (PIM and AIM) and Ctsk^+^Fmod^+^ PCs aggregated around the bone graft substitute had a close cellular interaction. PIM and AIM might influence the biological function of Ctsk^+^Fmod^+^ PCs through Spp1 signaling pathway.

### Characteristics of jaw bone Ctsk^+^Fmod^+^ PCs

To confirm the presence and location of Ctsk^+^Fmod^+^ PCs in rat jaw bone, we performed immunofluorescence staining at D3(fig. S7). Consistent with the ST results, there was a population of Ctsk^+^Fmod^+^ cells in the periosteum. We next attempted to isolate this population of cells from rat jaw bones. After getting single cell suspension, we labeled the cells with antibodies and acquired Lin^-^Ctsk^+^Fmod^+^ cells by FACS (fig. 4A). The sorted cells were a relatively pure group of Ctsk^+^Fmod^+^ cells by means of immunofluorescence staining(fig. 4B). The sorted cells had good colony-forming ability(fig. 4C). Additionally, Ctsk^+^Fmod^+^ cells could differentiate into osteoblasts *in vitro*(fig. 4D-E). We further compared the osteogenic potential of rat bone marrow-derived mesenchymal stromal cells (BMSCs) and Ctsk^+^Fmod^+^ PCs. ALP staining and quantitative analysis of ARS staining showed that Ctsk^+^Fmod^+^ PCs had osteogenic differentiation potential comparable to that of BMSCs(fig. 4D-E). Ctsk^+^Fmod^+^ PCs could also possessed the potential to differentiate into adipocytes and chondrocytes (fig. 4F-G). These data confirmed the existence of Ctsk^+^Fmod^+^ cells and validated its potential of multi-directional differentiation.

**Fig. 4.**
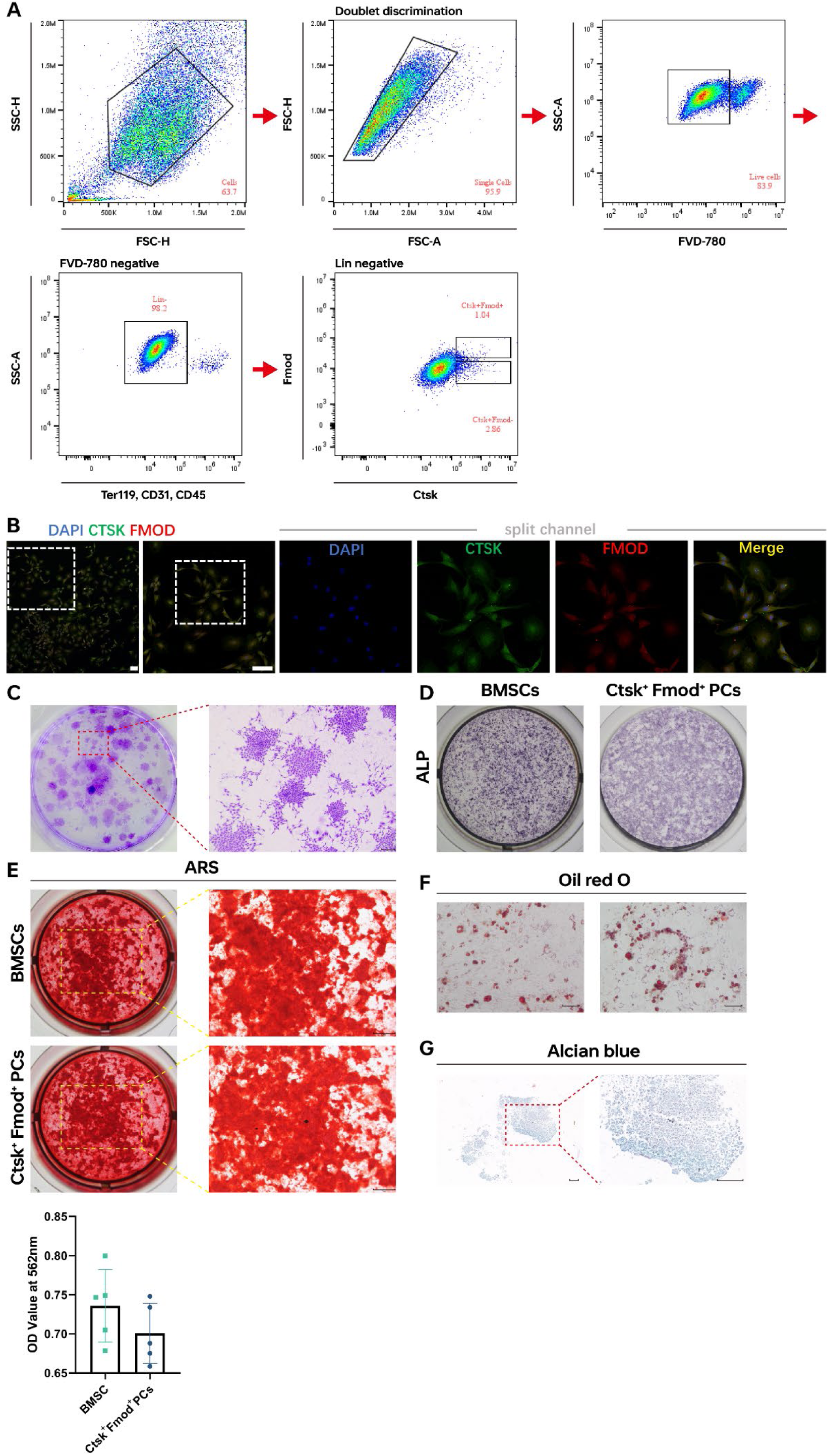
FACs analysis on rat periosteum and characteristic of Ctsk+Fmod+ PCs. (A) Schematic representation of the strategy used for FACs analysis of Ctsk^+^Fmod^+^ PCs. (B) Immunofluorescence results of sorted cells. Scale bar,100μm. (C) Colony formation of Ctsk^+^Fmod^+^ PCs at D14 via crystal violet staining. Scale bar, 500μm. (D) ALP staining of BMSC and Ctsk^+^Fmod^+^ PCs on D7. (E) ARS staining and quantitative analyses of BMSC and Ctsk^+^Fmod^+^ PCs on D21. Scale bra,1mm. n=5. (F) Oil red O stainining of Ctsk^+^Fmod^+^ PCs on D21. Scale bar, 100μm. (G) Alcian blue staining of Ctsk^+^Fmod^+^ PCs on D21. Scale bar, 100μm. ns, not significant by Student’s t-test.

### Spp1 secreted by macrophages reduced the osteogenic capacity of Ctsk^+^Fmod^+^ PCs

Notably, osteogenic related activities decreased in the Day7Bio group. We tried to understand what caused this phenomenon. According to the results of Cell-Chat and ST, we found that Spp1 signaling pathways were prominent between macrophages and Ctsk^+^Fmod^+^ cells in Bio group. In order to confirm the role of Spp1 secreted by macrophages, we co-cultured the rat macrophages and sorted cells via transwell system(fig. 5A). 48 hours later, we collected the supernatant from the upper chamber. ELISA results showed that macrophages in the co-culture system secreted more Spp1 protein(fig. S8A). This was in consistent with the PCR results that the *Spp1* gene expression of macrophages in the co-culture system was significantly increased(fig. S8B). In addition, a group supplemented with 2.0μg/mL mouse recombinant Spp1 protein(rmSpp1) were set up(*29*). ALP staining on Day7 showed that when adding rmSpp1 or co-cultured with macrophages, the osteogenic ability of Ctsk^+^Fmod^+^ PCs was decreased(fig. 5B). mRNA expression level of *Alp* and *Runx2* on Day7 was also down-regulated in co-culture group and this can be improved when blocking Spp1. (fig. 5C). ARS staining and quantitive analysis on D28 showed when Spp1 was blocked, the osteogenic effect of Ctsk^+^Fmod^+^ PCs was improved(fig. 5D).To further explore the effect of macrophages on Ctsk^+^Fmod^+^ PCs *in vivo*, we tried to achieve macrophage depletion in the surgical area by local injection of clodronate liposomes(*30*)(fig. 5E). The Immunofluorescence results of CD68 and flow-cytometric analysis showed the proportion of macrophages in the surgical area decreased after drug injection(fig. 5F, fig. S8C-D).Furthermore, the depletion of macrophages significantly reduced local *Spp1* gene expression in the surgical area. Histological examinations further demonstrated that new bone formation increased following the depletion of macrophages and downregulation of *Spp1*, according to the results of semi-quantitative analysis of new bone, new bone area, and expression of bone-related proteins ALP and OCN (fig. 5G-H, fig. S8E-F).

**Fig. 5.**
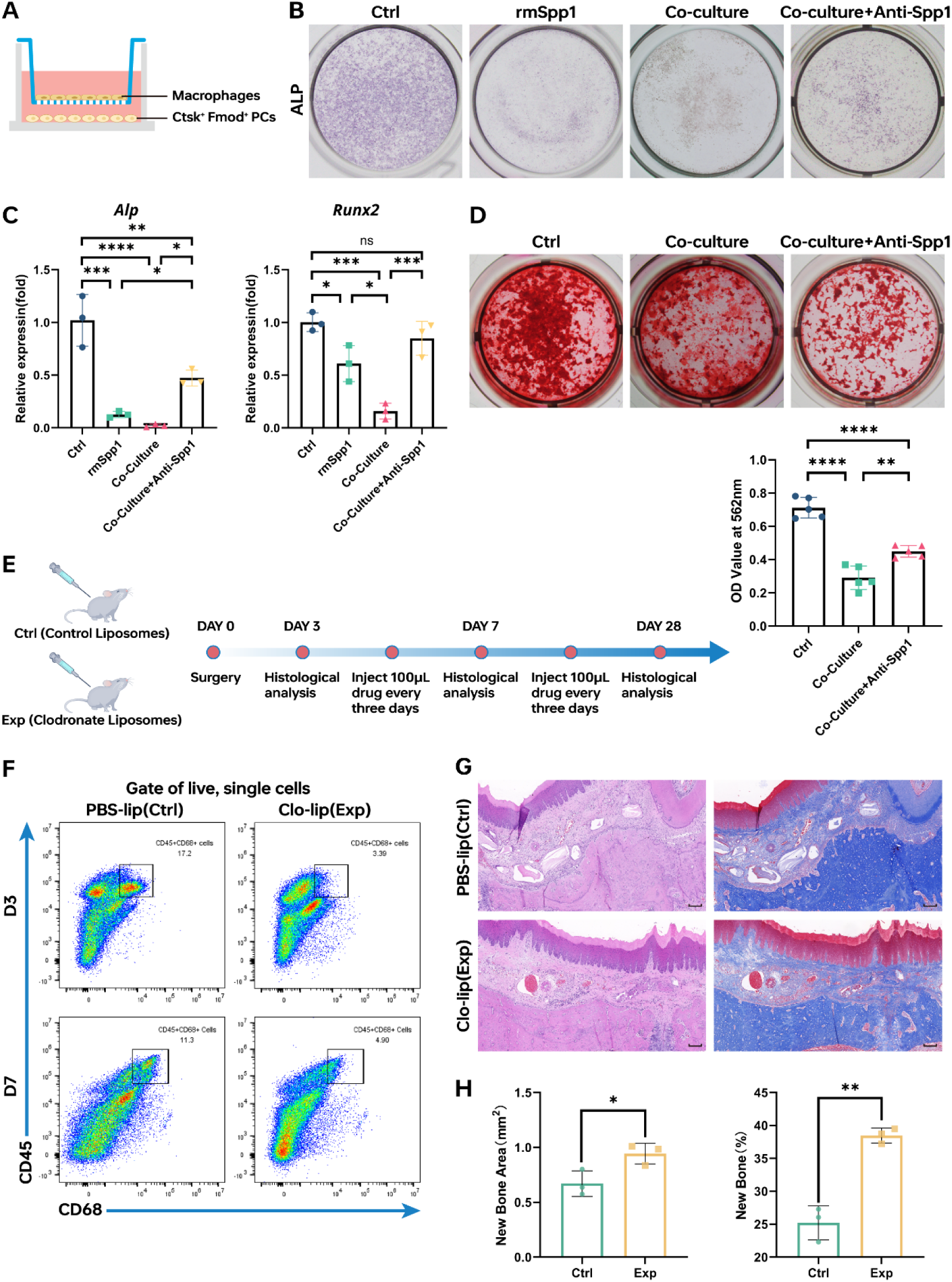
Spp1 secreted by macrophages reduced the osteogenic capacity of Ctsk^+^Fmod^+^ PCs. (A) Schematic representation of the co-culture system of macrophages and Ctsk^+^Fmod^+^ PCs. (B) ALP staining of PCs in different group on D7. (C)qPCR analyses of the mRNA levels of *Alp* and *Runx2* of Ctsk^+^Fmod^+^ PCs in different group. n=3. (D) ARS staining and quantitative analyses of PCs in different group on D21.n=5. (E) Workflow of macrophages depletion *in Vivo*. (F) *In vivo* administration of clodronate liposome depleted macrophages. (G) Histological staining of H&E and Masson after 28 days. Scale bar,100μm. (H) Quantitative analysis of new bone area and new bone. n=3. *P<0.05, **P<0.01,***P<0.001, **** P<0.0001, ns, not significant by one-way analysis of variance (ANOVA) with Tukey post-hoc test or Student’s t-test.

The above results all indicated that macrophages recruited to the surgical area secreted more Spp1, which impaired the osteogenic ability of Ctsk^+^Fmod^+^ PCs. Blocking Spp1 could increase the osteogenic potential of Ctsk^+^Fmod^+^ PCs and promoted osteogenesis in the surgical area.

## Discussion

Periosteum has the ability of bone regeneration and can be used for bone defect repair(*31*). Increasing evidence shows that functionally relevant cellular heterogeneity exist in the periosteum(*5, 6, 8, 10*). It’s a pity that most studies on the heterogeneity of periosteal cells are based on long bones. Because of the differences between long bones and jaw bones, the physical and cellular characteristics of periosteum differ according to the specific anatomical location(*11, 21, 32*). It’s important to analyze the characteristics of jaw bone periosteal cells for perfecting the theory of bone regeneration.

Combining the scRNA-seq and ST, we’re able to identify the key markers of PCs subsets and dissecting functional PCs subpopulations. Here, we constructed a subperiosteal bone grafting model inspired by clinical procedures(*20*). We mapped the rat PCs at single-cell resolution and identify the heterogeneity within the jaw bone PCs. PCs contained different cell subtypes with different functions, among which Ctsk^+^Fmod^+^ subset might be a population of cells having differentiation potential. Ctsk labeled a population of progenitors in the long bone and calvarial sutures(*5*). Periostin, highly expressed during development, plays an important role in PCs functions and fracture healing(*31, 33*). Thus we used Ctsk, Postn and Col3a1 to identify PCs for further analysis. Fmod, a coding gene of fibromodulin, is one of the small interstitial leucine-rich repeat proteoglycans (SLRPs) (*34*). Studies found that Fmod was more commonly distributed in oral soft tissues and helped to remain bone phenotype(*35, 36*). In this study, we found Ctsk and Fmod lable a subset of PCs. Combined with ST and immunofluorescence staining, we found that this population was located in the periosteal layer, and when periosteum was activated by the bone grafts, they would gradually migrate around the material. It was noteworthy that it was difficult to find the classic periosteal bilayer structure on day 7 in the Bio group, which may be related to the migration and remodeling of periosteal cells. By colony formation test and induced differentiation assay *in vitro*, we confirmed that Ctsk^+^Fmod^+^ PCs was a group of cells with good proliferation ability and differentiation potential. Ctsk^+^Fmod^+^ PCs that migrated around the bone grafts might differentiate into osteoblasts and participated in local osteogenic activities.

The bone graft implanted under the periosteum effectively activated the periosteum and provided space for bone formation. After implantation of the bone graft, a large number of macrophages were recruited. And we found that with a large number of macrophages infiltrating, local osteogenic activity was weakened. We nanlyzed the macrophage composition and found that PIM and AIM expressed higher Spp1 and interacted closely with Ctsk^+^Fmod^+^ PCs through Spp1(PIM and AIM)-Itga5/Itga8+Itgb1(Ctsk^+^Fmod^+^ PCs). One reason for the higher expression level of Spp1 was foreign body reaction (FBR). Researchers found that in FBR reaction, interactions of giant cells (GC) would be active and Spp1(GC)-Cd44(fibroblast) was prominent(*37*). However, we found that Spp1 expression in macrophages was also increased in the co-culture system. The effect of Ctsk^+^Fmod^+^ PCs on macrophages also contributed to the high expression of Spp1 in the Bio group. Spp1, also known as osteopontin, could influence the functions of cell adhesion, migration and survival with CD44(*38*).Spp1 is a key regulator of hematopoietic stem cell and plays as a physiologic-negative regulator of HSC proliferation(*39, 40*). Spp1 neutralisation could influence the response of liver progenitor cell and reduce fibrogenesis(*41*). However, the role of Spp1 in periosteal cells has not been elucidated. In our study, we found that Spp1 secreted by macrophages would impair the osteogenic ability of Ctsk^+^Fmod^+^ PCs. The expression of osteogenesis related genes (*Alp*, *Ocn*) was decreased, and the effect of osteogenesis was weakened, which could be partially rescued by blocking Spp1. When we reduced the proportion of local macrophages in the surgical area by injecting clodronate liposomes(*42*), the amount of new bone formation at day 28 increased.

In summary, this study uncovered the heterogeneity of PCs in jaw bone and identified Ctsk^+^Fmod^+^ subset which had multi-directional differentiation potential. Moreover, the interaction between PIM/AIM and Ctsk^+^Fmod^+^ PCs was explored. Spp1(PIM and AIM)-Itga5/Itga8+Itgb1(Ctsk^+^Fmod^+^ PCs) signalling impairs the osteogenic capacity of Ctsk^+^Fmod^+^ PCs, which could be partly rescued by blocking Spp1.

## Methods

### Animals

Wild-type, male SD mice purchased from Chengdu Dossy Experimental Animals Co., LTD. Were six-week-old and had an average weight of 200g. All the animal care and treatment procedures were carried out in accordance with international standards on animal welfare and approved by the Research Ethics Committee, West China Hospital of Stomatology, Sichuan University (WCHSIRB-D-2019-087).

### Subperiosteal bone grafting model

Briefly, after anesthesia, the hard palate mucosa was carefully elevated up with Dental Periosteal Elevator (Hu-Friedy) to ensure periosteum integrity. Next, Bio-Oss Collagen (Geistlich) was placed between the periosteum and hard palate as the experimental group (named Bio group) and a blank control group was set up. Finally, we suture the surgical area with cross-stitch using 7-0 suture (Prolene). After healing for 3, 7, 14 and 28 days, animals were euthanized for sample harvest.

### Sample processing, histology and histomorphometry

The harvested maxillae were fixed in 4% paraformaldehyde overnight at 4℃ and then stored in 75% ethanol for subsequent experiments. After fixed, the samples were decalcified in 10% EDTA for 4 weeks. Before embedded in paraffin, the samples were dehydrated through an ascending ethanol series. Longitudinal sections were obtained for H&E staining and Masson staining and the histology of the section was observed.

### Immunofluorescence and image analysis

The experimental procedure of immunofluorescence was consistent with our previous study(*24*). To determine osteogenesis and fibrogenesis in the subperiosteal region, we chose Alpl(Affinity, DF12525, 1:100) and Osteocalcin(Servicebio, GB11233, 1:1000). Immunofluorescence staining for Cathepsin K(Abcam, ab37259, 1:200)and Fibromodulin(Invitrogen, PA5-26250, 1:100) was performed for determination of the location of the PSCs.

### Single cell RNA sequencing

Specimen harvest procedure for scRNA-seq was reported in our previous reports(*24*). Briefly, we obtained three fresh samples from the surgical area per group(Bio and Ctrl) at different timepoints(Day3 and Day7), containing the hard palatal mucosa, periosteum, and the hard palate. We cut the specimen into 1mm width pieces and mixed them with type I collagenase (Gibco) and trypsin (Gibco) for sample digestion at 37℃ lasted for 2.5 hours. After dissociating, filtering, centrifuging the suspensions, we resuspended it in 3ml of red blood cell lysis buffer (Solarbio). Then we remove the dead cells and debris by Dead Cell Removal MicroBeads (Miltenyi). Last, 10x Genomics Chromium Single Cell 3′ v3 sequencing was performed using an Illumina 1.9 mode.

### Single-cell RNA-sequencing analysis

We used Cell Ranger pipeline software to generate the expression matrices for downstream analysis. After completing quality control and count normalization, we used the canonical correlation analysis for batch correction and integrated analysis (Seurat V4.1.2). Then we performed subpopulation analysis of PCs and macrophages. To infer the developmental progression of different subpopulation of PCs, we use the Monocle 2 package (V2.10.0) to order them in pseudotime. And we used Cellchat to predict communications among cell subpopulations.

### Spatial Transcriptions and spatial transcriptomic analysis

Fresh samples were harvested at different timepoints (Day3 and Day7) and groups (Bio and Ctrl). The samples were cryosectioned to get gene expression slides. After fixation, staining and imaging, the slides from both groups were ready for tissue permeabilization with the Visium Tissue Optimization Slide & Reagent kit. After reverse transcription, the spatially barcoded cDNA was released and collected for ST library preparation. The P5 and P7 primers, used in Illumina amplification, would be included in the final libraries.

We used SpaceRanger software to process raw FASTQ files and aligned histology images. The gene-spot matrices were analyzed with the Seurat package (V4.1.2). We used the SCTransform function to perform normalization across spots and independent component analysis (ICA) for dimensionality reduction. To generate the spatial cluster gene signature overlap correlation matrix, we analyzed all genes differentially expressed with average log2FC > 0.25 and adjusted p value < 0.05 across all ST clusters. Then we used the AddModuleScore function in Seurat to derive signature scoring from scRNA-seq and ST signatures. And we generated spatial feature expression plots with the SpatialFeaturePlot function.

### Fluorescence activated Cell Sorting (FACS) and cell cultures

To isolate Ctsk^+^Fmod^+^ PSCs, we obtained the periosteum from Jaw bones of 4-week-old SD rats (Chengdu Dossy Experimental Animals Co., LTD) and prepared single-cell suspension as described previously. The suspensions were incubated in the dark with primary antibody Cathepsin K (Abcam, ab37259, 1:300) and Fibromodulin (Invitrogen, PA5-26250, 1:300) for 1 hour. After being washed 2 to 3 times with 0.5% PB buffer, the cell solutions were co-incubated with FITC Goat Anti-Mouse IgG (H+L) Antibody (APExBIO, K1201,1:400) and APC Anti-Rabbit IgG (H+L) Antibody (Abcam, ab130805,1:400) for 0.5 hour. FACS was performed using BD FACS Ariall and analysis was performed using FlowJo 10.5.0.

Sorted cells were cultured in DMEM media(Gibco) with 20% FBS in 5% CO2 at 37℃. We replaced half of the media every 3 days and passaged cells once they were 70%-80% confluent.

### Colony formation test

500 sorted cells per well were seeded in 6-well plate. The medium was added as described above and it was changed every 3 days. After 10 days of culture, cells were fixed and stained with 1% crystal violet solution. Images were then taken with stereomicroscope (Olympus).

### Osteogenic, adipogenic and Chondrogenic differentiation

For osteogenic induction, sorted cells were expanded and then cultured in osteogenic differentiation medium with ascorbic acid (50 μg/mL), β-glycerophosphate (5 mmol/L), and dexamethasone (100 nmol/L). Media was changed every 2 days. 7 days later, ALP staining was performed with ALP staining kit (Beyotime).And after 21 days of induction, Alizarin red staining (ARS) were performed. 1% ARS Solution (Solarbio) was used to stain the mineralized nodules. Cells were then washed thoroughly for analysis.

For adipogenic induction, sorted cells were allowed to differentiation in adipogenic differentiation medium (Cyagen Biosciences, RAXMX-90031) and changed every 3-4 days for a total of 21 days. Being washed with PBS and fixed with 4% Paraformaldehyde, cells were stained with oil red O working solution and then observed under a light microscope.

For Chondrogneic differentiation, chondrogenic differentiation medium (Cyagen Biosciences, RAXMX-90041) was used in cell cultures. The media was changed every 3 days and allow to differentiation for 28 days. Then, after being washed with PBS and fixed, cells were stained with 1% Alcian blue solution.

### Isolation of rat macrophages

The macrophages were isolated from femur and tibia from 6-week rats. Briefly, the long bones were dissected and the bone marrow was flushed out. After centrifugation, the cells were resuspended in 5ml of red blood cell lysis buffer (Solarbio). Resuspended in α-MEM medium for more than 16 hours, the suspended cells were collected, centrifuged, and resuspended in α-MEM medium with 10% FBS, 1% penicillin/streptomycin and 50ng/mL M-CSF(Peprotech).

### Co-culturing of PSCs and rat macrophages

For transwell co-culturing, 2×10^4^ sorted cells were seeded into 24-well plate. The 0.4μm-pore size Corning transwell inserts(Sigma) containing 1000 macrophages were placed into the 24-well plate with sorted cells. The sorted cells were cultured in osteogenic differentiation medium and changed every 3 days. ALP staining and ARS staining were performed as described above at 7 days and 21 days later.

### Elisa

Spp1 levels were measured by collecting the supernatant from the upper chamber of the transwell co-culture system after 24h. The concentrations of Spp1 were determined according to the kit’s protocol (Jianglai Biotechnology).

### qPCR

Total RNA of PSCs and macrophages of the transwell co-culture system was purified with FastPure Cell/Tissue Total RNA Isolation Kit V2(Vazyme). cDNA was synthesized using HiScript III RT SuperMix for qPCR (Vazyme). The Taq Universal SYBR qPCR Master Mix (Vazyme) was used for qPCR on PCR instrument (qTOWER3G, Analytik Jena). Gapdh was used as an endogenous control and the 2^−ΔΔCt^ method was used to calculated the relative expression level of mRNA.

### Macrophage depletion in rat subperiosteal bone grafting model

Macrophages were depleted by local injection of 100μL clodronate liposomes(YEASEN) every 3 days. The ctrl group was injected with 100μL control Liposomes(YEASEN). After healing for 3, 7 and 28 days, animals were euthanized for sample harvest. H&E staining, Masson staining and immunofluorescence were performed when samples were fixed, decalcified and embedded in paraffin. The ImageJ software was used to analyze the results.

### Flow cytometry analysis

To confirm the success of macrophage depletion model, the fresh samples from the surgical area per group were obtained on day 3 and 7. After obtaining cell suspension by the above method, the cells were suspended in 300 μL staining buffer and incubated on ice for 30min with FVD780(eBioscience,1:1000) and then with CD45(Biolegend,202205, 1:300) for 30min. For intracellular antibody staining, cells were permeabilized and stained with CD68(Abcam, ab63856,1:5000) for 30min. The Beckman CytoFLEX Analyzer was used to test samples and Flowjo software (V10.8.1) was used to analyzed and visualized the flow cytometry data.

### Statistics

We dealt the data with Case Viewer software, ImageJ software, and GraphPad Prism 8.0 software. The statistical significance was analyzed using analysis of one-way analysis of variance (ANOVA) with Tukey post-hoc test or Student’s t-test. P < 0.05 was considered statistically significant(*P< 0.05, **P<0.01,***P<0.001, **** P<0.0001> and P > 0.05 was marked as ns (not significant).

## Acknowledgements

This work was supported by the This work was supported by the National Natural Science Foundation of China (grant number: 82271015, 81970965, 82201106).

---

**Fig. S1.**
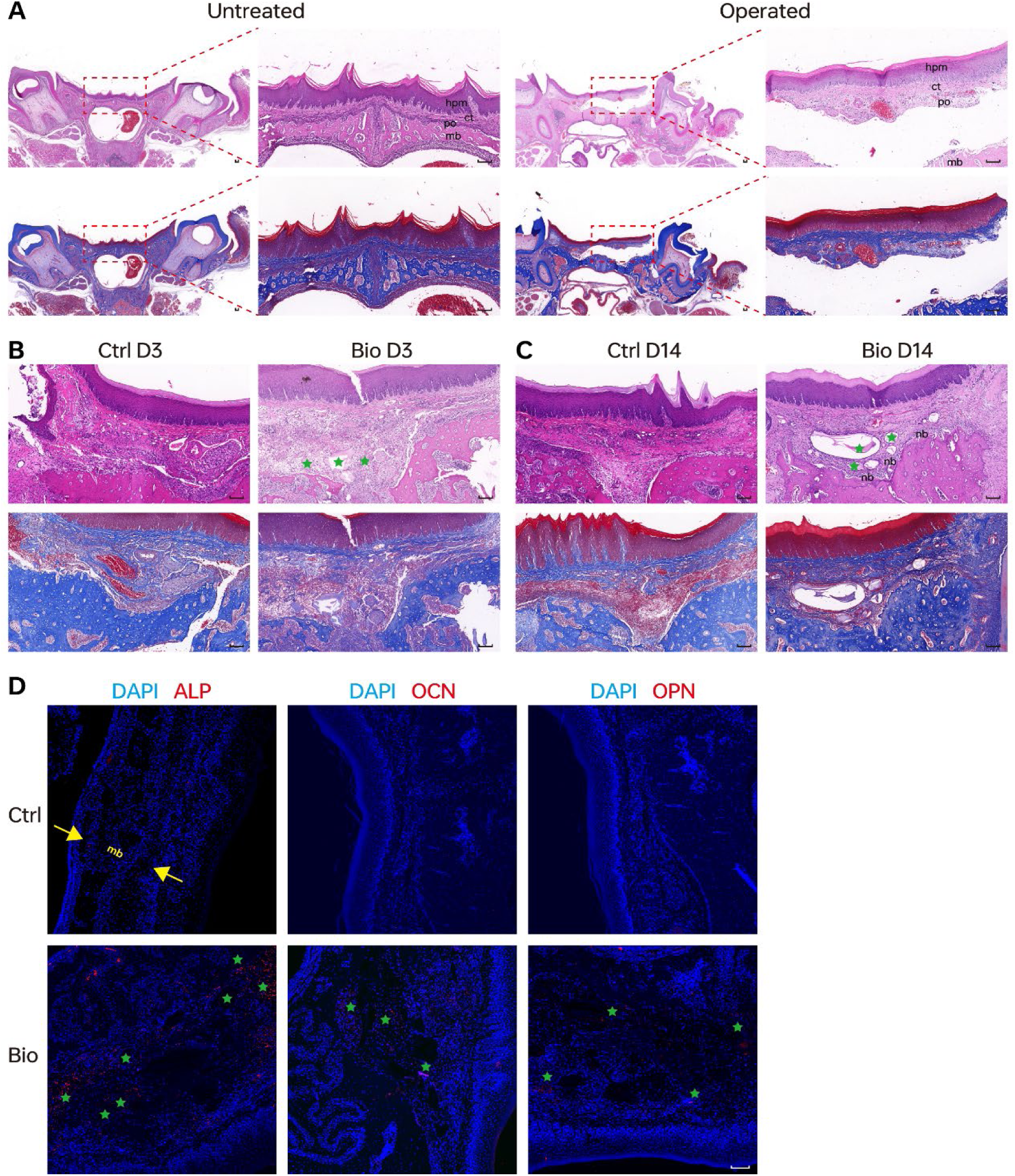
Histological analysis of rat subperiosteal bone grafting model. Representative H&E and Masson staining images of untreated and operated group(A) or Ctrl and Bio group on D3(B)、 D14(C). At post-surgical day 3, more blood cells and fibrin networks could be found in Bio group. New bone began to occur at D14 in Bio group near the bone graft. Scale nar,100 μ m. (D). Representative confocal images of sample sections from Ctrl and Bio group at D3. Notice the location of ALP, OCN and OPN expression. Scale nar,100 μ m. Blue stars, bone graft. Abbreviations: hpm, hard palate mucosa; ct, connective tissue; po, periosteum; mb, maxilla bone; nb, new bone.

**Fig. S2.**
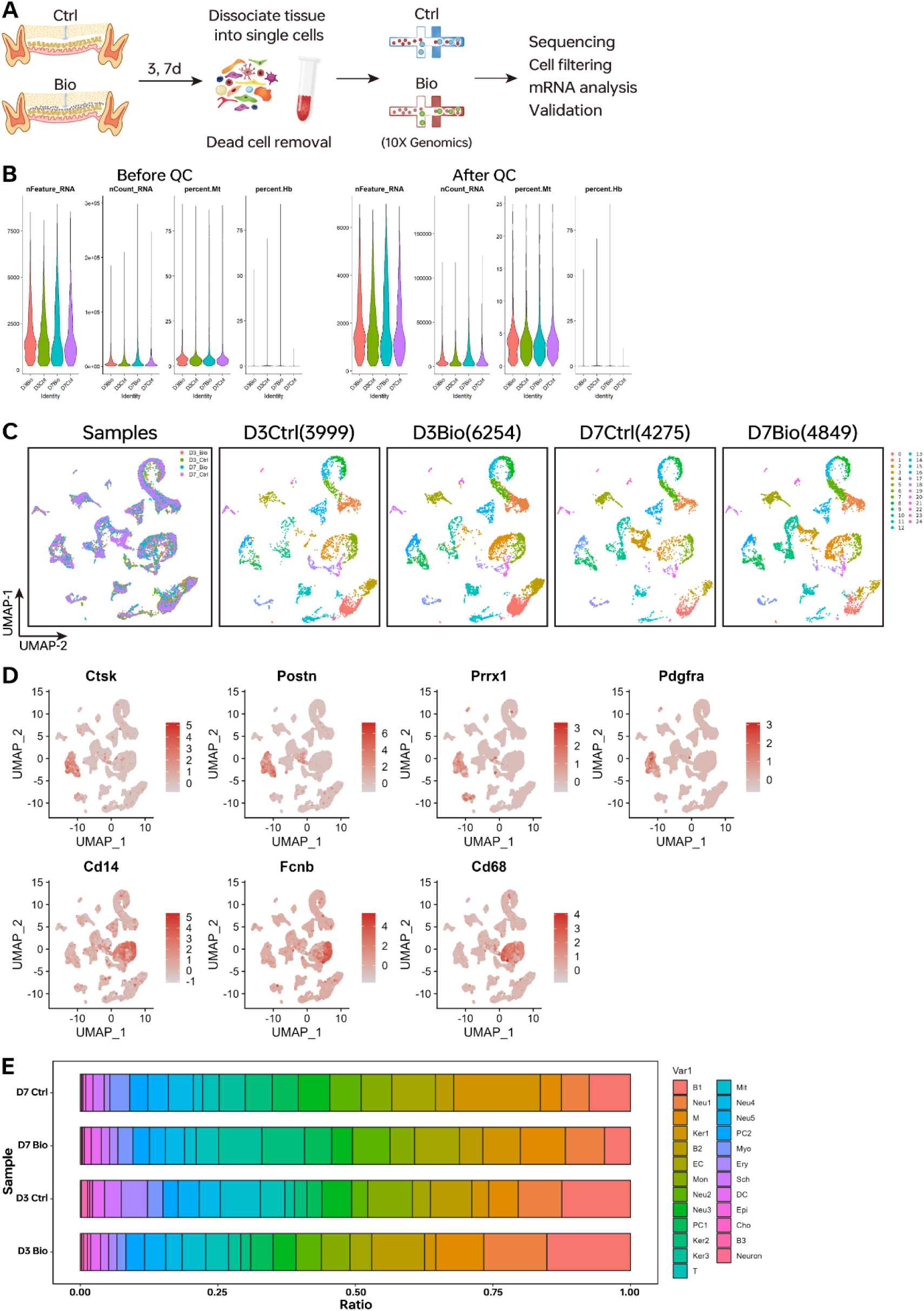
Overview of the scRNA-seq analysis between different group. (A) Workflow of scRNA-seq. (B) Violin plots showing the number of features, RNA counts, percent mitochondrial transcripts and percent hemoglobin in different group before and after quality control. (C) UMAP plots of cells at all time points(left) and at each time point of different group(right). (D) UMAP plots showing expression levels of selected known marker genes. (E) Bar plots showing proportion of all clusters in different group.

**Fig. S3.**
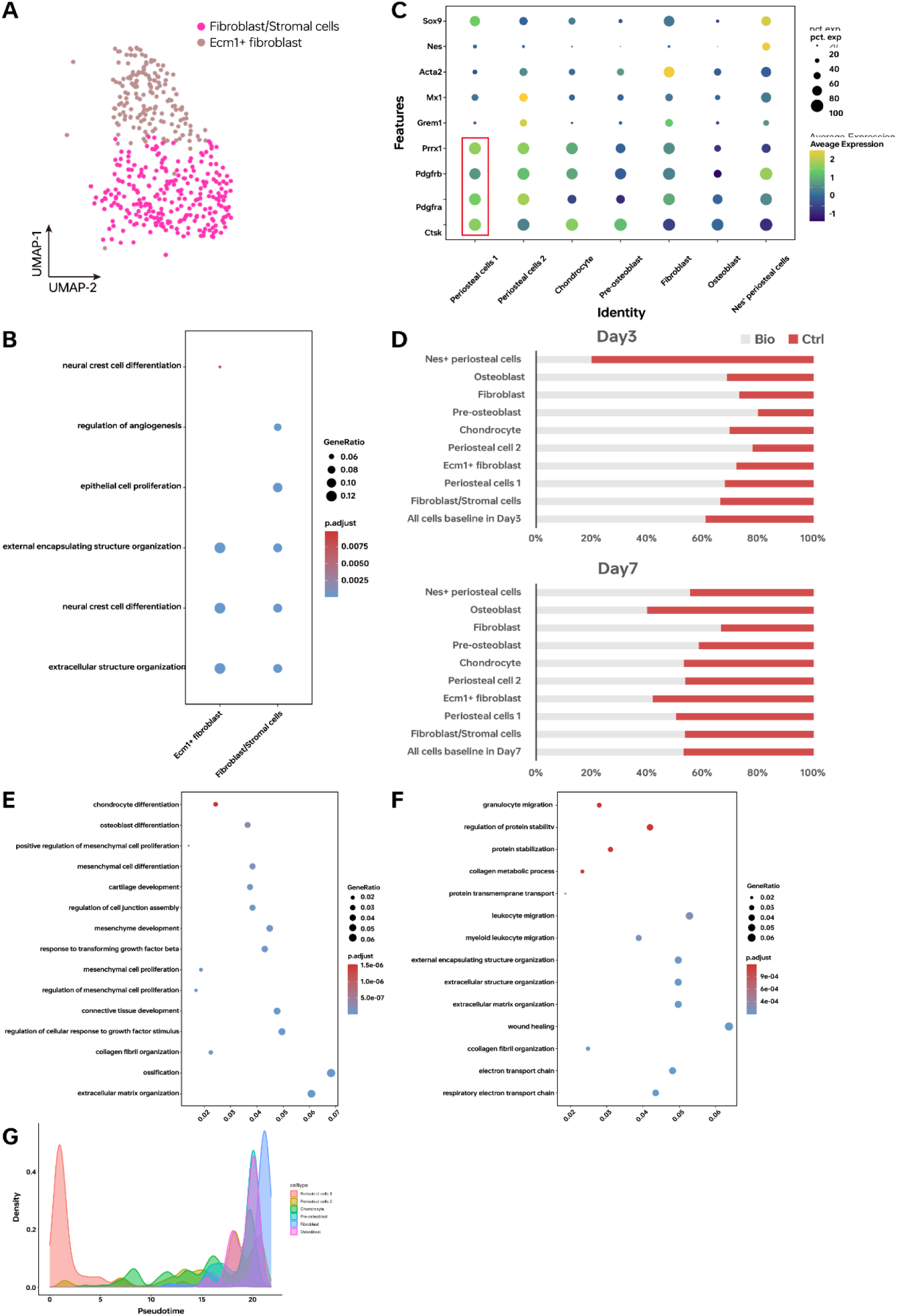
Further analysis of periosteal cells. (A) UMAP plots showing subclustering results of PCs. (B) GO analysis of subclusters of periosteal stromal cells. (C) Dot plots showing expression percent and average expression of stemness-related makers in subtypes of PCs. (D) Bar plots showing proportion of PCs-related cells in different group on Day3 and Day7. Differentially expressed genes (DEGs) of PCs between different groups on Day3(E) and Day7(F). (G) Plot of cell density over Pseudotime.

**Fig. S4.**
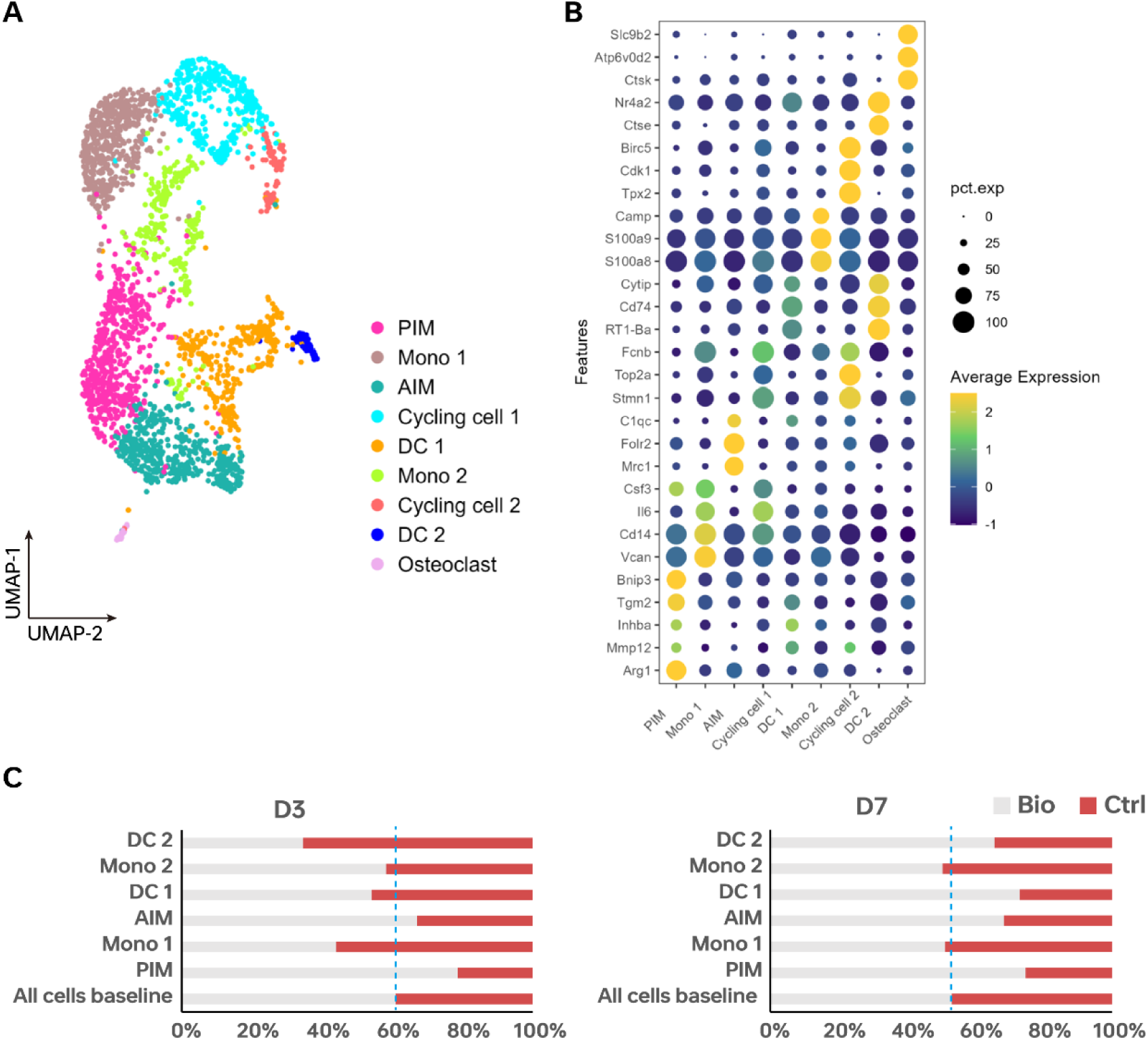
Subpopulation analysis of macrophages. (A) UMAP plots showing subclustering results of macrophages. (B) Dot plots showing expression percent and average expression of markers in subtypes of macrophages. (C) Bar plots showing proportion of macrophages subclusters in different group on Day3 and Day7.

**Fig. S5.**
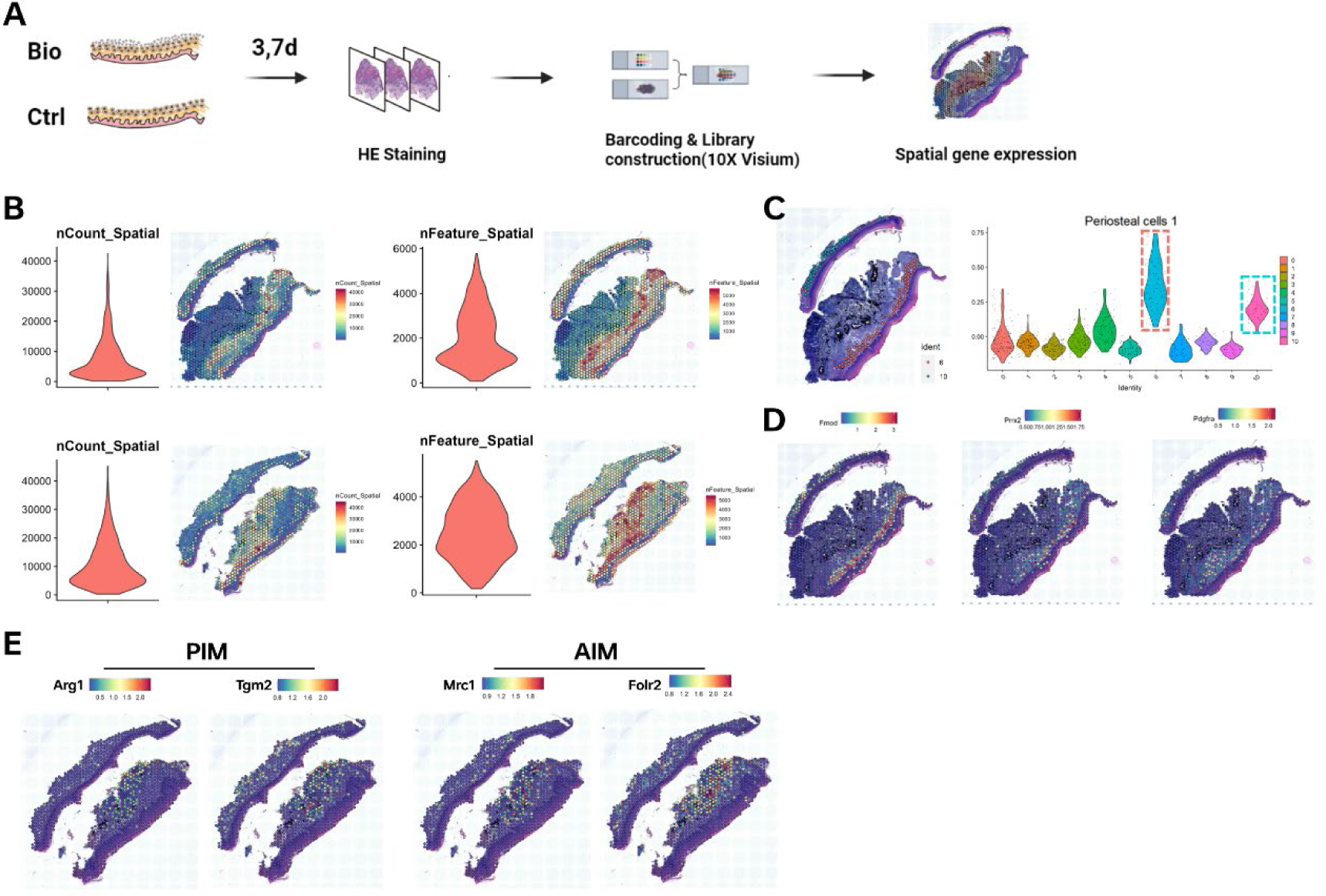
Spatial transcriptomics applied to Bio and Ctrl group. (A) Workflow of spatial transcriptomics (ST). (B) Violin plots of UMI counts and gene features per spot of different timepoints. (C) Spatial feature plots of periosteal cell 1-high clusters (Cluster 6 and 10) and Violin plots showing scores of its genes of individual spots derived from scRNA-seq for clusters of ST. Spatial feature plot highlighted the expression of marker genes for periosteal cell 1(D) and PIM or AIM(E).

**Fig. S6.**
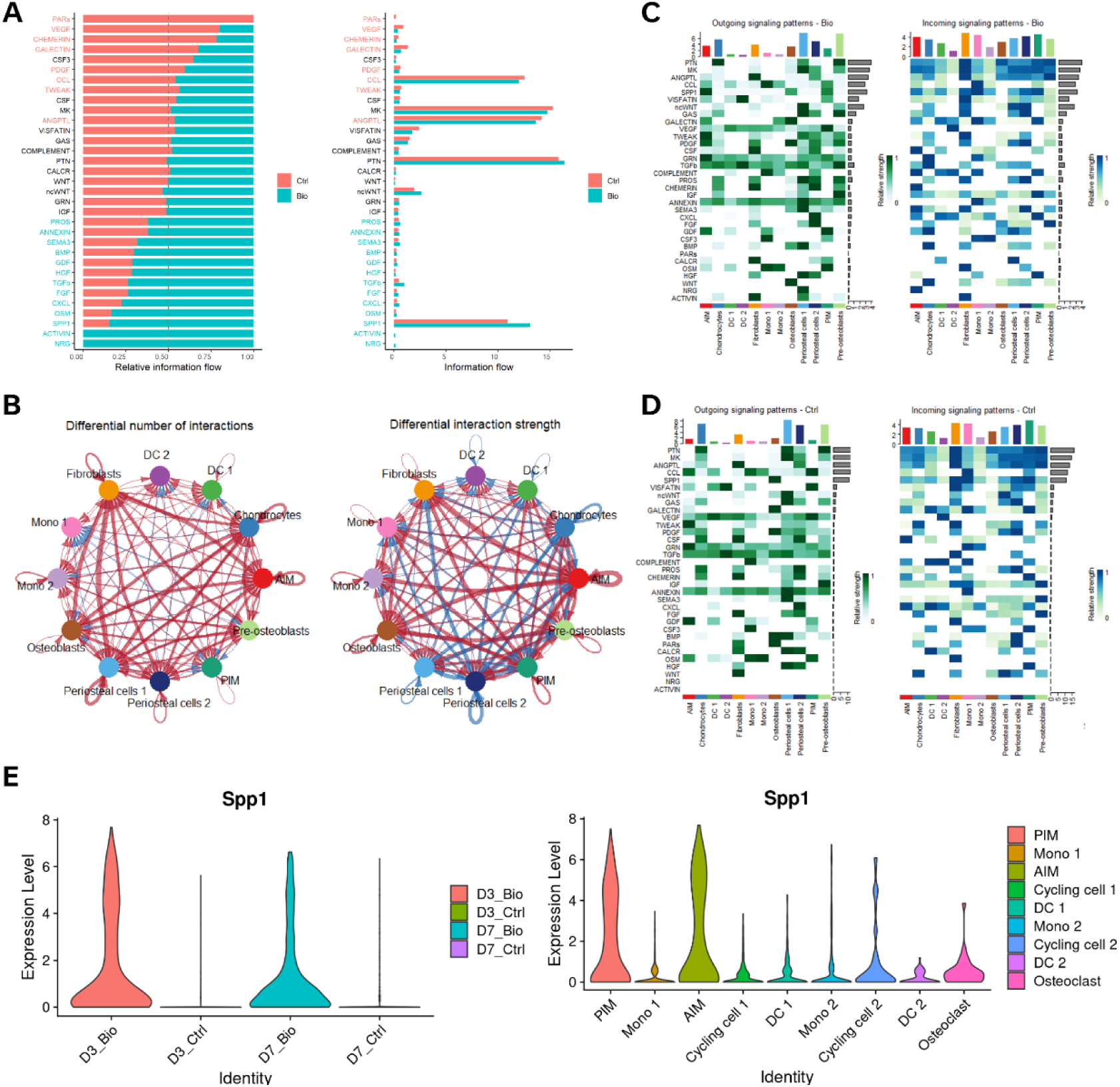
Cellular communication landscape between PCs and macrophages. (A) Comparison of the information flow between Bio and Ctrl group. (B) Net plot showing the interaction number and strength between each cluster. (C) Heatmap with the relative strength of each signaling pathways for clusters of PCs and macrophages in Bio group (C) and Ctrl group(D). (E) Violin plot for the expression levels of *Spp1* in different group or macrophage subsets.

**Fig. S7.**
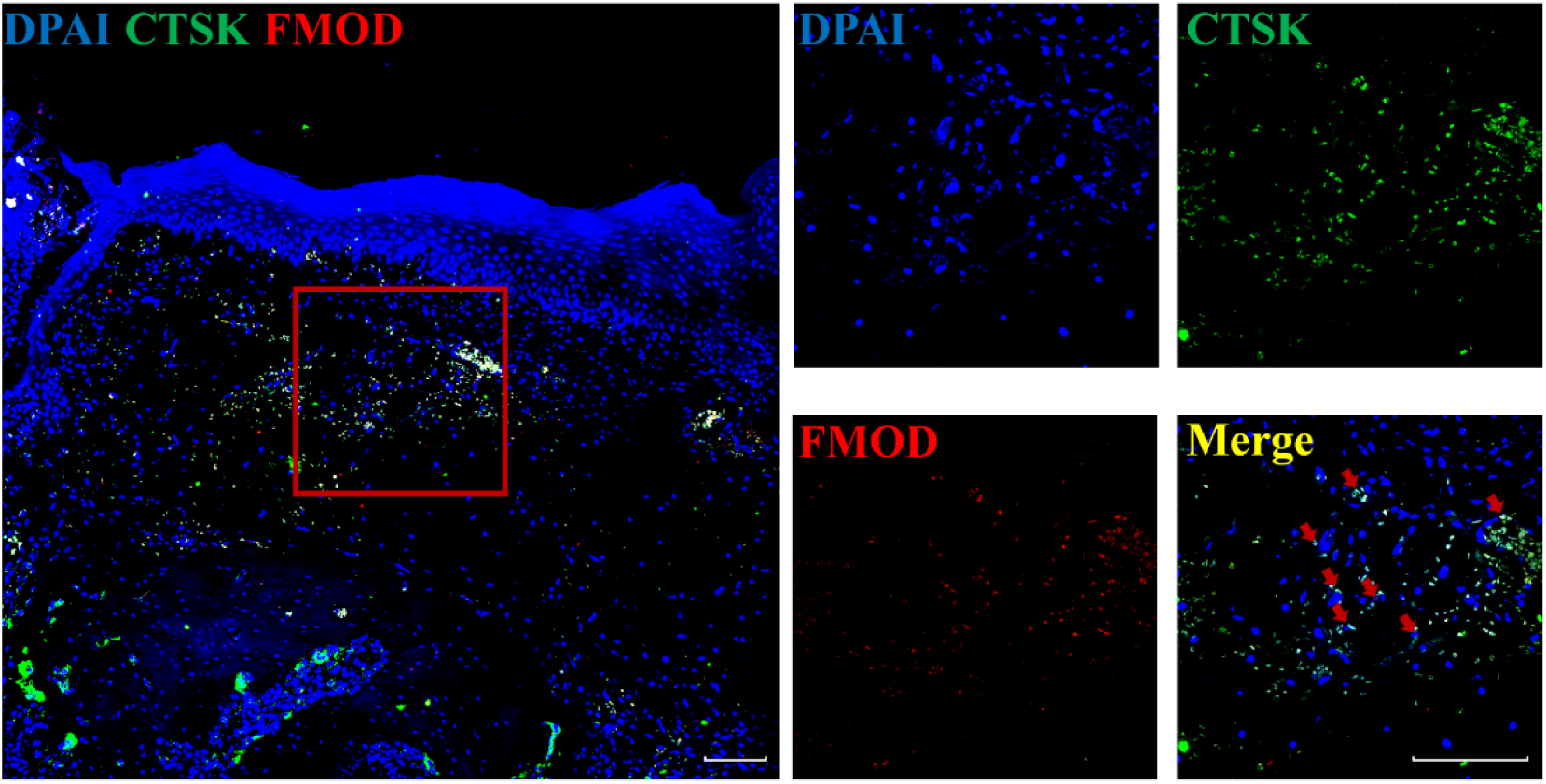
The presence of Ctsk^+^Fmod^+^PCs. Representative confocal images of sample sections from D3Bio group. The periosteum contained Ctsk^+^Fmod^+^ PCs, and their location was consistent with the results of ST. Red arrows, Ctsk^+^Fmod^+^ PCs. Scale bar, 100μm.

**Fig. S8.**
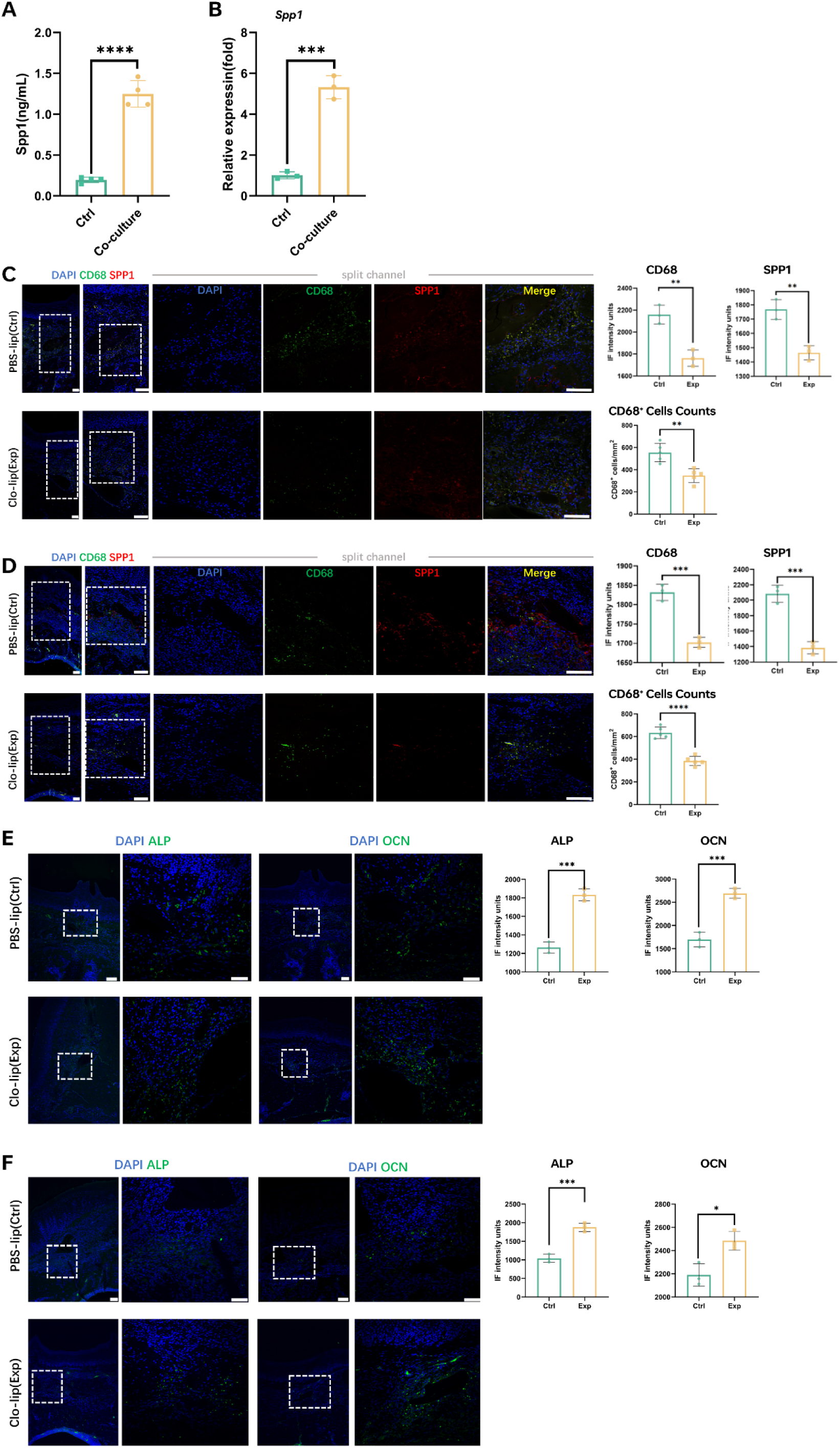
The role of Spp1 secreted by macrophages on Ctsk^+^Fmod^+^ PCs. (A) ELISA results of macrophage with or without co-culturing with Ctsk^+^Fmod^+^ PCs. (B) qPCR analyses of the mRNA levels of Spp1 in macrophages with or without co-culturing with Ctsk^+^Fmod^+^ PCs. Immunofluorescence staining of CD68 and Spp1 in operation areas at D3(C) and D7(D), or ALP and OCN in operation areas at D3(E) and D7(F), and normalized fluorescence intensity at D3 and D7. Scale bar, 100μm. *P<0.05, **P<0.01,***P<0.001, **** P<0.0001 by one-way analysis of variance (ANOVA) with Tukey post-hoc test or Student’s t-test.

